# Scanning SWATH acquisition enables high-throughput proteomics with chromatographic gradients as fast as 30 seconds

**DOI:** 10.1101/656793

**Authors:** Christoph B. Messner, Vadim Demichev, Nic Bloomfield, Matthew White, Marco Kreidl, Gordana Ivosev, Fras Wasim, Aleksej Zelezniak, Kathryn S. Lilley, Stephen Tate, Markus Ralser

## Abstract

Bridging genotype to phenotype, the proteome has increasingly become of major importance to generate large, longitudinal sample series for data-driven biology and personalized medicine. Major improvements in laboratory automation, chromatography and software have increased the scale and precision of proteomics. So far missing are however mass spectrometric acquisition techniques that could deal with very fast chromatographic gradients. Here we present scanning SWATH, a data-independent acquisition (DIA) method, in which the DIA-typical stepwise windowed acquisition is replaced by a continuous movement of the precursor isolation window. Scanning SWATH accelerates the duty cycles to a few hundreds of milliseconds, and enables precursor mass assignment to the MS2 fragment traces for improving true positive precursor identification in fast proteome experiments. In combination with 800 µL/min high-flow chromatography, we report the quantification of 270 precursors per second, increasing the precursor identifications by 70% or more compared to previous methods. Scanning SWATH quantified 1,410 Human protein groups in conjunction with chromatographic gradients as fast as 30 seconds, 2,250 with 60-second gradients, and 4,586 in conjunction with 5-minute gradients. At high quantitative precision, our method hence increases the proteomic throughput to hundreds of samples per day per mass spectrometer. Scanning SWATH hence enables a broad range of new proteomic applications that depend on large numbers of cheap yet quantification precise proteomes.

## Introduction

Proteomes bridge between genotype and phenotype, and are likewise important for basic as well as data-driven biology, as they are for biotechnology and systems medicine^1–4^. There is an increasing need to record proteomes at large numbers and at high quality. Indeed, there are several applications, like functional drug screens, or epidemiological scale human studies, that are restrained by the throughput of proteomics. Proteomes, however, are inherently complex, generating huge analytical challenges to be recorded in a short amount of time, and, to maintain high quantification precision if they are acquired at large numbers^5^.

Several recent developments have addressed this problem and increased sample throughput and quantification precision at the steps of sample preparation and data analysis. Automatization and 96-well plate based sample processing allow the preparation of hundreds of samples per day and reduce batch effects, that otherwise are a challenge in large-scale and longitudinal experiments^6–12^. Further, fast, efficient and robust chromatographic separations have been achieved by replacing nanoflow LC, as traditionally used in proteomics ^13,14^, with setups that use higher flow-rates. This ranges from microflow LC systems (5-50µl/min)^15–17^, to novel LC devices with preformed gradients ^18,19^. More recently, we have introduced proteome experiments that make use of high-flow liquid chromatography (800 µl/min). In 5 minute chromatographic gradients, these allowed up to 180 proteome injections/day on a single LC-MS instrument, while increasing robustness, cost-efficacy and quantification precision in longitudinal proteome experiments ^10^. In addition, the development of algorithms that enable the efficient deconvolution of complex spectra as resulting from fast chromatographic measurements is still an ongoing process, but several major steps have been achieved recently, and have increased proteomic depth as well as quantification precision in conjunction with the fast chromatographic experiments ^20–23^.

Still missing are however mass spectrometric acquisition methods that are specifically designed for the challenges of high-throughput proteomic experiments. Proteomic experiments that make use of the desirable short chromatographic gradients (i) require a high sampling velocity in the chromatographic dimension. For instance in 5-minute high-flow chromatography proteomic experiments, peaks can elute at a full width of half maximum (FWHM) of 3 seconds or less ^10^. Moreover, (ii) separating complex samples in short gradients produces complex spectra with a high degree of signal interferences that occur when samples are separated with lower peak capacities. Indeed, ion trap-type mass spectrometers that have been widely used in the proteomic field have been predominantly operated using data-dependent acquisition (DDA) schemes that directly depend on the sampling velocity of the mass spectrometer. DDA techniques achieve a high dynamic range and provide high-quality identification and quantification data ^1^. DDA methods, however, become limited in depth and consistency, when the number of co-eluting peptides exceeds the sampling velocity. A popular solution to this problem has hence been to use long chromatographic gradients ^24^, or to decrease the sample complexity via 2-dimensional chromatography ^25,26^ or pre-fractionation ^27^. While providing excellent data quality and depth in small scale experiments, this strategy results in long measurement times and increases batch effects in large measurement series. An alternative to DDA approaches is to replace the selection of individual precursor ions with the sampling of wide mass windows, giving rise to data-independent acquisition (DIA) approaches ^28^, such as MS^E 29,30^ or SWATH-MS^31^. In SWATH-MS, the mass spectrometer is configured for stepping through a predefined set of wide precursor isolation windows, thus consistently fragmenting all the precursors within the mass range of interest^31^. This way SWATH-MS reduces the undersampling problem and can increase the identification numbers and consistency in the analysis of complex samples in single-shot proteomics^32,33^. However, SWATH-MS is also limited when proteomic experiments are done with fast chromatographic gradients. SWATH-MS with fast chromatographic gradients requires short duty cycle times. These can be achieved by reducing the number of isolation windows, but as a consequence these become wider, which results in co-fragmentation of co-eluting precursors, reducing depth ^23^.

To address the challenges of fast proteomic experiments, we have developed a DIA method as well as adequate software. In scanning SWATH, the stepwise windowed acquisition of SWATH-MS (Figure 1a) is replaced by continuous scanning with the first quadrupole, using a quadrupole Time-of-Flight (qTOF) mass spectrometer (TripleTOF6600 ^34^, Sciex). The scans allowed us to accelerate the acquisition duty cycles to a minimum of 280 milliseconds, enabling us to record proteomes with sub-minute chromatographic gradients. Further, scanning SWATH adds a novel dimension to the DIA data due to the sliding mass window, that captures the fragments in a time-dependent as well as precursor mass-dependent manner. We have advanced our software, DIA-NN ^23^, to exploit this dimension by assigning precursor masses to MS2 fragment traces, which increase true positive peptide identifications in the complex convoluted spectra produced by the fast scanning SWATH experiments. We demonstrate that the combination of scanning SWATH and industry-standard high-flow chromatography (800 µL/min) allowed us to conduct high-quality proteome experiments with gradient lengths as fast as 30 seconds and that we achieve comprehensive proteomic depth with gradients as fast as a few minutes, hence substantially augmenting the high-throughput capacity of proteomics.

**Figure 1:**
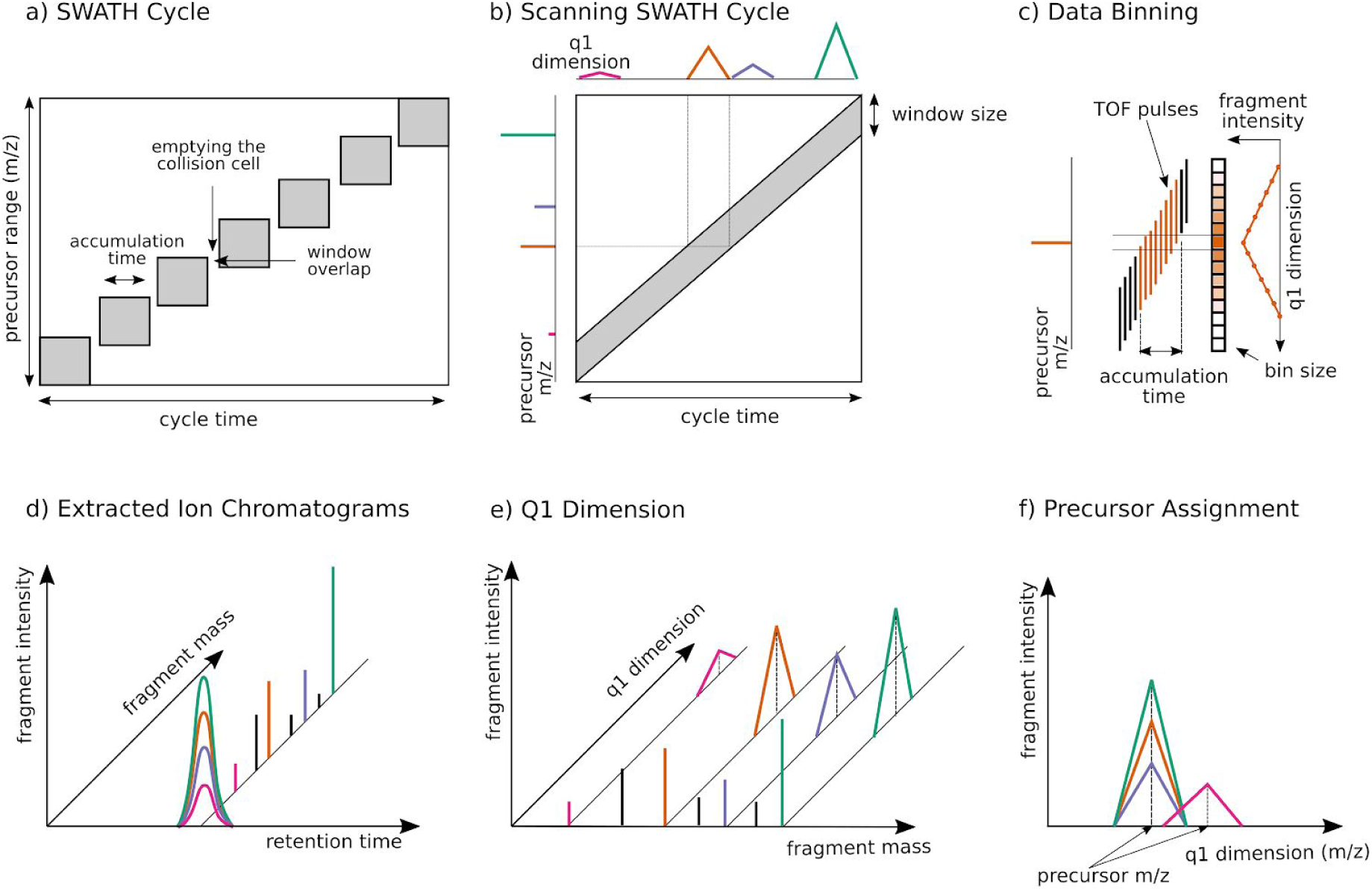
Scanning SWATH replaces the stepwise precursor selection with a continuously moving quadrupole and thereby adds another dimension to the data: **a**. In conventional SWATH-MS/DIA-MS, a quadrupole selects a relatively wide mass range and the detector collects MS/MS spectra for a defined accumulation time. The individual chromatograms used for quantification are then reconstructed computationally and post-acquisition. The windows are stepped and are overlapping (to compensate for edge effects^35^). The collision cell needs to be emptied after each step. **b**. In scanning SWATH, the isolation window slides over the precursor mass range and MS/MS spectra are continuously acquired. The cycle time is only defined by the scan speed and not by the window size. The continuous movement of the quadrupole results in a time dependency of the fragment intensity. Fragment signals appear when the leading edge of the quadrupole passes the precursor m/z and they disappear when the precursor m/z falls out of the quadrupole isolation window. In this dimension, the MS2 signal intensities are time-dependent, with the maximum at the precursor m/z. This added dimension can hence be exploited to assign precursor masses to MS2 fragment traces (as in ***e***). **c**. The acquired raw data is sectioned into bins of a defined m/z size. Data from TOF pulses that overlap with a certain m/z bin are summed together and written into the respective bin (e.g. all TOF pulses labeled in red on the diagram are summed together in the respective bin). Therefore, the highest signal for a certain fragment is in the bin which includes the precursor mass. However, in contrast to conventional SWATH, data from each TOF pulse is written into more than one bin, resulting in a Q1 profile of a triangular shape. **d**. The q1 profile provides the 4^th^ dimension of scanning SWATH data. As in conventional SWATH, each fragment mass (mass dimension) has a certain intensity (intensity dimension) that is measured along the chromatographic time (retention time dimension). **e**. In scanning SWATH data, each precursor is assigned to a q1-profile (q1 dimension) **f**. Different fragments from the same precursor show correlating q1 profiles and the center of the q1 profile corresponds to the precursor mass.

## Results

### Replacing the stepped precursor isolation with continuous scans accelerates the duty cycle and adds a scanning dimension to DIA data

In conventional SWATH-MS, precursor ions are selected for fragmentation via cycling stepwise through a predefined set of isolation windows (Figure 1a)^31^. For precise quantification, an acceptable number of data points per peak (ideally, 5-7 ^33^) are required. In long chromatographic gradients conventional to proteomics (peak width > 10 seconds) such is achieved with window sizes between m/z 15 and m/z 30, to reach duty cycles in the range of 2 - 4 seconds^35^.

In scanning SWATH, we achieve duty cycles starting from 200 milliseconds, enabling us to record a similar amount of points per peak in conjunction with much faster chromatographic gradients. In scanning SWATH, precursor ions are selected using a “sliding” isolation window, which moves across the m/z range of interest, with fragmentation spectra being continuously recorded by the TOF analyzer (Figure 1b and 1c). These scans can be completed in a shorter time than conventional stepped SWATH acquisitions as there is no need to empty the collision cell between steps. Further, the cycle time does not depend on the window size as in conventional SWATH. Scanning SWATH, therefore, “decouples” cycle time and window size and allows to run fast duty cycles with narrow isolation windows.

In order to determine an ideal window size for the identification and quantification performance, we ran yeast (*S. cerevisiae*) whole proteome tryptic digests on a 5-minute high-flow water-to-acetonitrile gradient ^10^. We recorded the data using scanning SWATH with a fast cycle time of 500 ms and window sizes ranging from 3 m/z to 20 m/z, covering a precursor range from 400 m/z to 900 m/z. The best results in terms of identifications and quantitative precision were achieved with window sizes as narrow as 10 m/z (Figure S1a). Reducing the window size further would result in even higher identification numbers due to less interference but the resulting shorter effective accumulation times would lower the quantitative precision. Therefore, we set the window size to 10 m/z for all further experiments.

### The scanning quadrupole enables the assignment of precursor masses to fragment traces which reduces the number of false identifications

So far, a disadvantage of SWATH-MS over DDA acquisition techniques has been a lack of precursor mass assignment to the MS2 traces ^35^. The continuous movement of the quadrupole in scanning SWATH results in a time dependency of the fragment signal and adds a further dimension to scanning SWATH data. The signal of each MS2 feature first appears and then disappears when the leading margin and the trailing margin of the “sliding” isolation window pass the precursor mass, respectively (Figure 1b). The acquired data is written into defined m/z bins by summing up all TOF pulses that overlap with the respective precursor range (Figure 1c). The resulting triangular “Q1 profiles” can be mapped to m/z coordinates by calibrating on known masses and aligning the Q1 profiles with the respective MS1 mass (see Methods). We introduced changes in our open-source software DIA-NN^23^ that allow exploiting this dimension to assign precursor masses to each MS2 feature observed. This dimension is complementary to the retention time dimension as it allows to distinguish co-eluting peptides with different precursor masses (Figure 1, lower panel).

In order to test to which extend scanning SWATH and the use of the Q1 dimension improves identifications in short gradients, we injected 10 µg of a trypsin digested Human cell line (K562) proteome and ran a 5-minute high-flow chromatographic gradient. Using a Q1 sliding window size of 10 m/z, the data was written into 2 m/z bins, providing a resolution in the Q1 dimension that allows the effective use of the Q1 scores. As false-discovery rate (FDR) calculations are software- and acquisition mode-specific, which might affect benchmarking results, we compared scanning SWATH data to conventional stepped SWATH using the two-species library approach, which estimates true positive calls in an unbiased fashion on the basis of an experimentally measured FDR ^23,36^. To do this, we augmented a Human spectral library with *Arabidopsis thaliana* precursors as negative controls and used them to calculate FDR based on the ratio of known true and false positives (i.e. the calling of an *Arabidopsis* specific peptide in the Human sample would be considered a false positive,^23,36^, Methods). In the five minute gradient, scanning SWATH identifies 70% more true positive precursors at 1% FDR than a highly optimized, stepped SWATH acquisition method ^10^ (Figure 2a). This improvement originates from a combination of the fast duty cycle and the ability to identify false positives via the matching of the precursor mass to MS2 features. For example, for the true target (human precursor-AVVIVDDR(2+)), the apex of the Q1 profile matches the mass of the precursor in the library and thus increases the confidence in this particular identification (Figure 2b - left panel). On the other hand, the apex of the Q1 profiles that correspond to the extracted fragment masses of a false target (*Arabidopsis thaliana* precursor-FDGALNVDVTEFQTNLVPYPR(3+)) do not match their respective precursor mass (Figure 2b-right panel). Therefore, this particular false target had a reported Q-value that was above 0.01 (not identified) but was incorrectly identified in the conventional SWATH run (reported Q-value below 0.01). Thus, the use of the Q1 profile allows to increase the number of identifications by better distinguishing true targets from interferences.

**Figure 2:**
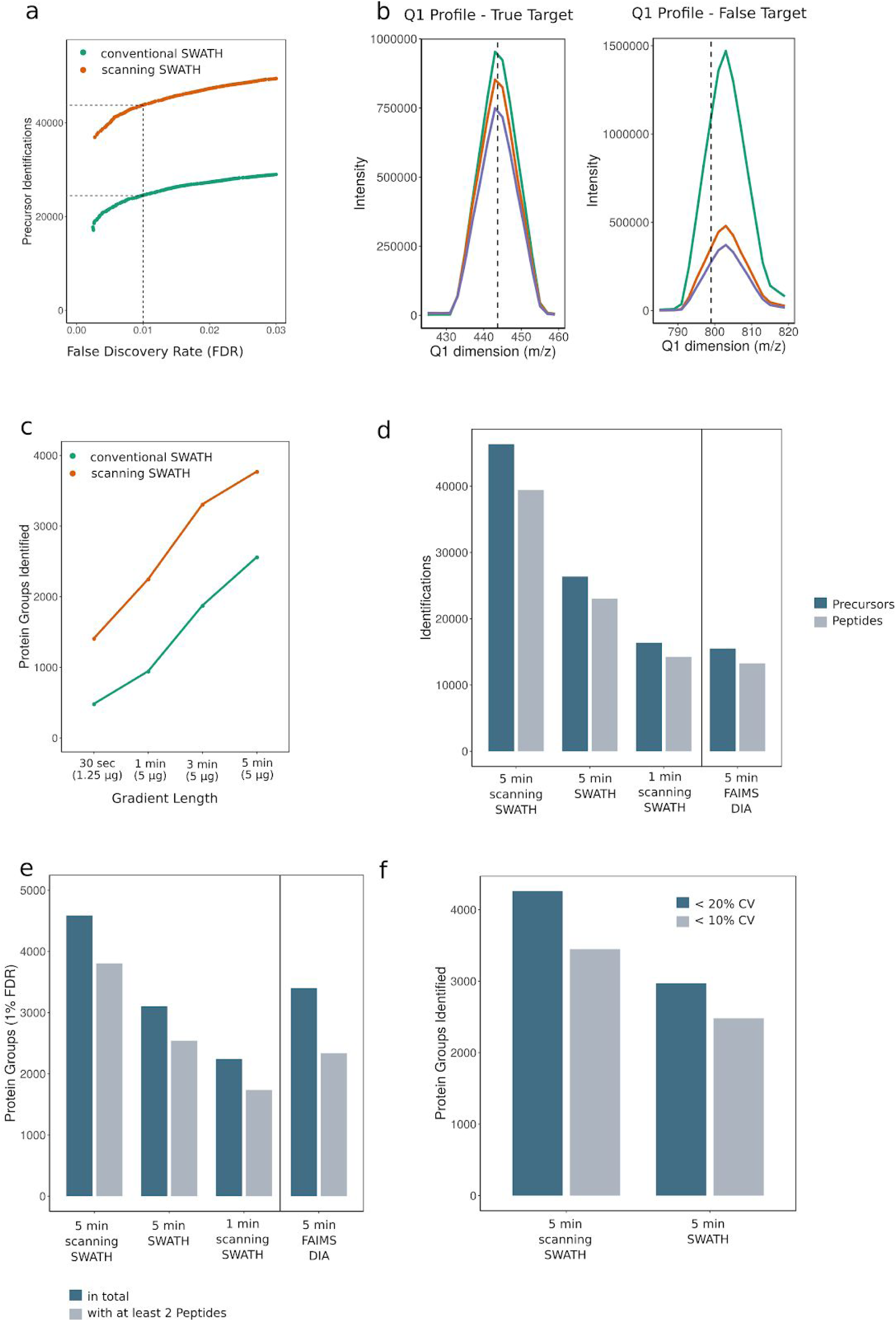
Scanning SWATH enables sub-minute proteomics, and increases peptide identifications in fast high-flow chromatographic gradients. **a**. Scanning SWATH improves protein precursor identification in fast high-flow-chromatographic experiments by >70% compared to conventional SWATH. Scanning SWATH (10 m/z window) and conventional stepped SWATH (based on an optimized method^10^) runs (5-minute water to acetonitrile chromatographic gradient, 800µL/min) of 10 µg K562 cell line tryptic digests were analyzed with a two-species library, that contains Human and *Arabidopsis thaliana* precursors. Number of identified Human precursors is shown as a function of the experimentally estimated false discovery rate (FDR)^23,36^. **b. left panel:** Q1 profile of fragments corresponding to a true target precursor (Human) with the mass of 443.8 m/z (AVVIVDDR(2+)). The three most intense fragments are shown (716.4 m/z; 617.3 m/z; 504.2 m/z)). The center of the q1 profiles of the fragments match the precursor mass. **b. right panel:** Q1 profile of MS2 features corresponding to the fragment masses of a false target (*Arabidopsis thaliana* precursor) with the mass 799.1 m/z (FDGALNVDVTEFQTNLVPYPR(3+)). This precursor had the reported Q-value above 0.01 (not identified) in the scanning SWATH run but was incorrectly identified in the conventional stepped SWATH run (reported Q-value below 0.01). **c**. Number of protein groups identified in a Human cell lysate (K562) with scanning SWATH and conventional stepped SWATH, using 5, 3, 1 minute and 30 second linear 800 µL/min chromatographic gradients. The amount of proteins injected was 5 µg for the 5,3 and 1-minute gradients and 1.25 µg for the 30-second gradient. The duty cycles were adjusted to the gradient length (Table S2,S3 and S4). I.e they were as fast as 280 milliseconds for the 30-second gradient. **d**. Number of precursors (peptides ionized to a specific charge) and peptides (stripped sequences) identified (1% FDR) in Human cell lysates measured with different acquisition schemes and platforms. A Human cell lysate (K562) was acquired with five and one-minute gradient scanning SWATH (“1 min scanning SWATH” and “5 min scanning SWATH”) and 5-minute conventional stepped SWATH (“5 min SWATH”) on a high-flow UPLC (Agilent Infinity II) coupled to a q-TOF (SCIEX TripleTOF 6600) instrument. 10 µg human cell lysate (K562) was injected for the 5-minute gradients and 5 µg for the 1-minute gradient. The normal SWATH method is based on a short gradient SWATH method published recently^10^, with slightly modified cycle time and precursor range to match the parameters of the scanning SWATH method, to allow direct comparison (see Methods). The data was further compared to recently published 5-minute gradient DIA runs of cell lysates (HeLa) acquired on Orbitrap Exploris 480 with a FAIMS coupled to an Evosep One system (“5min DIA FAIMS”) (PXD016662)^19^. In order to make the identification numbers comparable, the runs were analyzed with project-specific libraries generated on the respective setups using DIA-NN (see Methods). Identification numbers are the mean of triplicate injections for “5 min scanning SWATH”, “5 min SWATH” and “5 min DIA FAIMS”. **e**. Number of protein groups with at least one or two peptide identifications, respectively. **f**. Number of protein groups quantified with a coefficient of variation (CV) below 20% and below 10% in triplicate injections with the different acquisition schemes and platforms.

### Scanning SWATH records quantification-precise proteomes in combination with chromatographic gradients as fast as 30 seconds

An ideal technology to achieve fast gradients is high-flow chromatography, with flow rates of several hundred microliters per minute, and short column lengths, that reduce the washing and equilibration cycles between the runs. We have recently shown that short gradient high-flow chromatography offers further benefits to proteomic experiments, because of a high peak capacity, longitudinal chromatographic stability, and the stability of high-flow electrospray ^10^. Furthermore, we achieved a proteomic throughput of 180 samples/day, including all washing and equilibration steps, that were in the range of just 3 minutes ^10^. Here we make further use of core-shell particles as the stationary phase (Methods), which enable low back pressures and highly efficient separations ^37,38^. To test to which extend scanning SWATH enables proteome experiments with fast high-flow gradients, we ran linear gradients with 5, 3, and 1 minute as well as 30-second length at a flow rate of 800 µL/min and acquired the data with correspondingly adjusted duty cycles (see Methods), for a Human cell line tryptic digest (K562). With a scanning SWATH cycle time of 280 ms (Table S2), we were able to record sufficient data points per peak (3 average at FWHM) in conjunction with the 30-second chromatographic gradient. Processing the data with DIA-NN, the 30-second gradient quantified 1,410 protein groups at 1% FDR. The identification numbers increased to 2250 protein groups with a 60-second chromatographic gradient (cycle time of 310 ms), and to 3772 protein groups (1% FDR) in five minutes (cycle time of 520 ms) when 5 µg Human cell lysate were injected (Figure 2c).

To put the performance of scanning SWATH into the context of other methods, we illustrate a benchmark in which we compare the 5-minute scanning SWATH runs (“5 min scanning SWATH”) with 1-minute scanning SWATH (“1 min scanning SWATH”), with 5-minute conventional stepped SWATH (same LC-MS setup) (“5 min SWATH”), and with recently published DIA data recorded with an Orbitrap Exploris 480 instrument (Thermo) with FAIMS interface and with 5-minute separations on an Evosep One (“5 min DIA FAIMS”) ^19^. The latter is to our knowledge the only publicly available proteomics DIA dataset with gradients as fast as 5-minutes and thus puts our data into context of a complementary technology. We note that the DIA-FAIMS runs applied different MS parameters. For example, the precursor range and cycle time (1 sec) differ from our MS settings. Further, the Evosep One LC system used has different chromatographic properties with lower flow rates and makes use of single-sample filter tips ^39^. In 5-minute gradients and with 10 µg human cell lysate injected, scanning SWATH achieves an increase of more than 70% in precursor identifications (peptides ionized to a specific charge) compared to stepped SWATH on the same setup (46,342 vs 26,372) (Figure 2d). A 60 seconds gradient scanning SWATH run identifies 16,373 precursors which is more than the number of precursors identified with the 5-min Evosep-DIA-FAIMS runs ^19^(15,533) if analyzed with the same software and software settings (Methods) (Figure 2d). Further, scanning SWATH identifies 4,586 protein groups (4,016 unique Proteins (only proteotypic peptides considered)) while conventional stepped SWATH and DIA-FAIMS identified with the same gradient length 3,102 and 3,400 protein groups, respectively (Figure 2e). Out of these 3,804, 2,539 and 2,336 are identified with at least 2 peptides in 5-minute scanning SWATH, 5-minute conventional SWATH and 5-minute DIA-FAIMS, respectively.

Despite the increase in the identification numbers, scanning SWATH maintains a high quantification precision with a median coefficient of variation (CV) value of 4.9% for all proteins quantified. Scanning SWATH shows a higher precision than conventional SWATH (Figure S1b), with median CV values of 3.5% and 4.2% for scanning SWATH and conventional SWATH, respectively, when comparing on the same set of proteins (proteins quantified in both, scanning as well as conventional SWATH runs). Also in absolute numbers scanning SWATH performs significantly better, as it quantifies 4,261 protein groups (out of 4,586 identified) with < 20% CV and 3,449 with < 10% CV, while conventional SWATH quantifies 2,971 and 2,481 with < 20% and < 10% CV, respectively (Figure 2e). The precision values obtained with the Evosep-DIA-FAIMS ^19^ emerge from the use of the different acquisition method with slower cycle times, chromatographic device and single-sample filter tips for each injection, and are substantially lower (Figure S1b).

## Discussion

Here we demonstrate the acquisition of precise proteomes in fast gradients, as enabled by scanning SWATH, a data-independent acquisition technique. This method accelerates mass spectrometric duty cycles for proteome experiments to less than 300 milliseconds, while allowing narrow precursor isolation windows, and adds an additional scanning dimension to the raw data. Through additions to our software DIA-NN^23^, we exploit this scanning dimension to assign precursor masses to MS2 fragments for improving true positive peptide identifications out of complex data resulting from very short chromatographic gradients. We report, to our knowledge for the first time, the quantification of more 1,410 protein groups in conjunction with a chromatographic gradient as fast as 30 seconds, and show that with slightly longer gradients (60 seconds to 5 minutes), at least 70% more precursors are quantified compared to previous DIA methods, or compared to alternative high-throughput DIA-methods. Despite this high speed, the quantification precision is comparable if not better to the most recent achievements (Figure 2f). By quantifying the median protein group with a precision of CV < 4.9%, indicate that despite the high-throughput, the combination of high-flow chromatography and scanning SWATH is among the most precise proteomic methods currently available.

By using high-flow (800 µL/min) chromatography scanning SWATH allows measuring several hundreds of proteomes per day on a single LC-MS instrument (i.e. 180 samples per day in conjunction with the slowest of the employed gradients (5 minutes), including a ∼3 minute overhead between runs for washing, equilibration and sample loading), while the stability of the high-flow rate chromatographic regime helps to reduce batch effects and to increase measurement quality for large-scale projects ^10^. We benchmarked the platform with Human cell lysates, due to its high complexity, but it is equally applicable to sample types with less complexity (e.g. yeast or plasma/serum samples), where one can make maximum use of the fast throughput. The amount of proteins injected was 1.25 µg to 10 µg, which is an accessible protein amount with conventional digestion protocols^10,40–42^. For instance, the digestion of just 5µL of blood plasma would allow five to ten injections on our platform. These injection amounts should not be misinterpreted as scanning SWATH being less sensitive; the injection amounts are a consequence of the intentional choice to use the high-flow chromatography as an ideal setting for achieving the fast gradients and stable high throughput experiments ^10^. For other applications than the one intended in our study, i.e. those that have low sample amounts available, scanning SWATH can equally be coupled to conventional proteomic nanoflow or microflow chromatography, and profit from their high sensitivity on low sample amounts. Indeed, also microflow chromatography has recently been adapted to achieve decent throughput in proteomic experiments ^16,17^. Finally, we would like to discuss that scanning SWATH requires a fast qTOF mass spectrometer, but does otherwise not depend on expensive or proprietary reagents, and all of our software is freely available and easy to use (i.e. DIA-NN contains an intuitive graphical interface, and runs on both Windows as well as Unix based operating systems ^23^). As open source software, our approaches to analyse scanning SWATH data can be incorporated in both academic as well as commercial software developments. In addition to boosting throughput and measurement precision, high-flow-scanning SWATH does hence reduce costs. For instance, in combination with our recently presented sample preparation workflow optimized to achieve ISO-level certification, the material costs for each proteome are below 10$ ^10^, and the total costs are even much lower if using sample preparation methods that avoid the use of expensive SPE plates, like the SP3 method ^7^. In combination, these developments render high-throughput proteomics faster and significantly cheaper than many other omic techniques.

The ability to run more than a thousand low-cost proteomes per week per mass spectrometer enables a series of new applications, ranging from precision medicine, over systems biology, to quality control applications. For instance, we have recently shown that quantitative assessment of plasma proteomes could classify COVID-19 patients at a time, where the immune evasion strategies of the SARS-CoV-2 virus were still barely understood ^10^. Indeed, our choice of analytical standard high-flow chromatography reduces the burden to implement proteome technologies in clinical laboratories. Second, the new high throughput capacities render proteomics applicable to drug screens, were due to a lack of sufficient throughput, so far molecular readouts were largely restricted to transcriptional profiling or metabolomics ^43,44^.

## Methods

### Materials

Water (LC-MS Grade, Optima; 10509404), Acetonitrile (LC-MS Grade, Optima; 10001334) and Formic acid (LC-MS Grade, Thermo Scientific Pierce; 13454279) were purchased from Fisher Chemicals. Human cell lysate (MS Compatible Human Protein Extract, Digest, V6951) were purchased from Promega.

### Sample preparation

The Human cell lysate was obtained commercially (Promega) and the yeast digest was prepared as previously described^23^. The digested peptides were dissolved in 3% Acetonitrile/0.1% Formic acid.

### Liquid chromatography - mass spectrometry

Liquid chromatography was performed on an Agilent Infinity II ultra-high-pressure system coupled to a Sciex TripleTOF 6600. The peptides were separated in reversed-phase mode using a InfinityLab Poroshell 120 EC-C18, 2.1 × 50 mm, 1.9 μm particles column, and a column temperature of 30°C. If not mentioned otherwise, a gradient was applied which ramps from 3% B to 36% B in 5 min (Buffer A: 1% acetonitrile and 0.1% formic acid; Buffer B: acetonitrile and 0.1% formic acid) with a flow rate of 800 µL/min. For washing the column, the organic solvent was increased to 80% B in 0.5 min and was kept for 0.2 min at this composition before going back to 3% B. An IonDrive Turbo V Source was used with ion source gas 1 (nebulizer gas), ion source gas 2 (heater gas) and curtain gas set to 50, 40 and 25. The source temperature was set to 450 and the ionspray voltage to 5500V.

For comparing different gradient length (0.5, 1, 3 and 5 minutes) we applied linear gradients ramping from 3% B to 36% B (Buffer A: 1% acetonitrile and 0.1% formic acid; Buffer B: acetonitrile and 0.1% formic acid) with a flow rate of 800 µL/min. We injected 5 µg for the 5, 3, and 1-minute gradients and 1.25 µg for the 30-second gradients. For the scanning SWATH and conventional stepped SWATH the duty cycles were adjusted accordingly (Table S2 and S4). For conventional SWATH this was done by adjusting the number of variable windows to reach cycle times comparable to the scanning SWATH duty cycles (Table S3 and S4). For this particular comparison, the accumulation times of the MS1 scan was 10ms and of the MS2 scans 25ms.

### Scanning SWATH operation

The scanning SWATH runs were acquired with a scanning SWATH beta version. If not mentioned otherwise the following settings were applied in the scanning SWATH runs: The precursor isolation window was set to 10 m/z, a mass range from m/z 400 to m/z 900 was covered in 0.5s and the raw data was binned in the quadrupole or precursor dimension into 2 m/z bins. The MS1 scan was omitted and the data was acquired in high sensitivity mode with a total protein amount of 10 µg injected.

The instrument control software calculates an RF/DC ramp which is applied to quadrupole filter 1. The ramp is calculated from the experiment start transmission mass, stop transmission mass, transmission width, and cycle time. The calculation uses previously acquired calibrations to calculate ramps for mass DACS and resolution DACS. The quadrupole start mass is calculated as experiment start mass minus transmission width, and, the quadrupole stop mass is experiment stop mass plus transmission width. This allows for correct precursor profiles of all fragments at the boundaries of the experimental mass range. Collision energy is calculated using the +2 Rolling Collision energy equation based on the center masses for each transmission window. This results in a small collision energy spread depending on the width of the transmission window relative to the range being scanned. In these experiments the effect is typical around 1 eV spread for a given precursor.

Scanning SWATH calibration is automated when running a pre-built batch and directly infusing a tuning solution (ESI Positive Calibration Solution for the SCIEX X500 System (SCIEX)) with the 266.16; 354.21; 422.26; 609.28;829.54) Quadrupole response of each standard is measured at transmission window widths 3,5,10,15,and 20 m/z where each width is additionally measured scanning at 500, 1000, 2000 and 3000 m/z / sec. The recorded quadrupole responses for each condition are stored in a three-dimensional matrix. The dimensions are width, speed and m/z. The values stored in the matrix are observed m/z from theoretical m/z. Observed precursor m/z is calculated from current pulse number relative to total scan pulses applied as a fraction of scanned mass range plus start mass. An exact calibration curve is therefore known for each of the acquired scan speeds and widths. For scan speeds and widths in between resulting from experimental parameters, a curve is tri-linearly interpolated.

The instrument acquisition software organizes ion detection responses into calculated 2 m/z precursor isolation bins given the current ToF pusher pulse number relative to the start of the scan applying the scanning SWATH offset curve described above. The 2 m/z precursor isolation bins are organized in the data file as adjacent experiments allowing for the extraction of precursor profiles for any given fragment in a given cycle by tracing fragment response across experiments as well as normal chromatographic profiles across cycles.

### Conventional DIA and SWATH runs (for benchmark)

The conventional 5-minute “stepped” SWATH method is based on a previously published method^10^. To make it comparable to the developed scanning SWATH method we applied the same 0.5-second duty cycle and the same precursor mass range from m/z 400 to m/z 900 as in the developed scanning SWATH method. Each duty cycle consists of one MS1 scan with 50ms accumulation time and 17 MS2 scans with variable windows (Table S1) and 25ms accumulation time.

The DIA-FAIMS data acquired on an Evosep One LC system coupled to an Orbitrap Exploris 480 was downloaded from ProteomeXchange (dataset PXD016662). Triplicate runs with 500ng HeLa tryptic digests loaded on column (highest load in this dataset), a compensation value of -45 V for FAIMS, a resolving power of 15,000 and a cycle time with 1s were considered as this runs provided the best identification numbers while maintaining quantitative accuracy ^19^. The DIA-FAIMS data was analyzed with a project-specific library acquired on the same setup (PXD016662, “5min-library.kit”). For the analysis in DIA-NN, the library was exported from Spectronaut (v. 13.12.200217.43655 (Laika)) with the “Export Spectral Library” function and reannotated with the “Reannotate” function in DIA-NN using the UniProt^45^ human canonical proteome (3AUP000005640). The DIA-FAIMS data was analyzed with Spectronaut (v. 13.12.200217.43655 (Laika)) and DIA-NN but as the identification numbers were higher with DIA-NN we used these values for the benchmark.

### Data analysis

Raw data processing was carried out with DIA-NN (Version 1.7.11) with default settings in “robust LC (high accuracy)” mode. Plots were generated with R ^46^. For calculating the protein CV values, protein quantities were obtained using the MaxLFQ algorithm ^47^ as implemented in the diann R package (https://github.com/vdemichev/diann-rpackage).

Extracted ion chromatograms were generated with the PeakView software (Version 2.2, SCIEX).

### Matching precursors to MS2 fragment traces in DIA-NN

DIA-NN takes full advantage of the 4^th^ dimension in scanning SWATH data. In DIA-NN, a set of scores is calculated for each precursor-spectrum match (PSM), to distinguish true signals from noise using linear classifiers and an ensemble of deep neural networks. DIA-NN also selects the ‘best’ fragment ion per PSM, as the one with the clearest signal, with other fragment ions then being assessed by comparing their MS2 traces to those of the best fragment^23^. Scores specifically related to Q1 profile assessment have now been added to DIA-NN algorithms. Briefly, a Q1 profile (as illustrated in Figure 1c) is extracted (at the putative peptide elution peak apex) from the MS2-level data for each fragment ion m/z, as well as for the non-fragmented precursor ion trace. DIA-NN then calculates scores that reflect (i) how similar are the Q1 profiles of the fragments and the non-fragmented precursor to the Q1 profile of the best fragment, (ii) how well the Q1 profile shapes match the expected triangular shape, and (iii) how well the centroid of the Q1 profile corresponding to the best fragment matches the precursor mass. All the scores are calculated in triplicate: using either 3, 7, or 11 bins closest to the putative Q1 profile apex.

### Library generation

The libraries were generated from “gas-phase fractionation” runs using scanning SWATH and small precursor isolation windows. 5 µg of K562 cell lysate (Promega) or 5 µg yeast digests were injected and run on a nanoAcquity UPLC (Waters) coupled to a SCIEX TripleTOF 6600 with a DuoSpray Turbo V source. The peptides were separated on a Waters HSS T3 column (150mm x 300µm, 1.8µm particles) with a column temperature of 35°C and a flow rate of 5 µL/min. A 55-minute linear gradient ramping from 3% ACN/0.1FA to 40% ACN/0.1% FA was applied. The ion source gas 1 (nebulizer gas), ion source gas 2 (heater gas), and curtain gas set to 15, 20 and 25 respectively. The source temperature was set to 75 and the ion spray voltage to 5500V. In total 11 injections were run with the following mass ranges: m/z 400-450, m/z 445-500, m/z 495 - 550, m/z 545-600, m/z 595-650, m/z 645 - 700, m/z 695 - 750, m/z 745 - 800, m/z 795- 850, m/z 845 - 900, m/z 895 - 1000 and m/z 995 - 1200. The precursor isolation window was set to m/z 1 except for the mass ranges m/z 895 - 1000 and m/z 995 - 1200, where the precursor windows were set to m/z 2 and m/z 3, respectively. The cycle time was 3sec consisting of high and low energy scan and data was acquired in “high resolution” mode. A spectral library was generated using library-free analysis with DIA-NN directly from these scanning SWATH acquisitions. For this DIA-NN analysis, MS2 and MS1 mass accuracies were set to 20 ppm, respectively, and scan window size set to 6.

### Empirical FDR estimation with two-species library

In order to empirically validate the FDR we generated a two-species library as described previously^23,36^. Briefly, we augmented the Human library with *Arabidopsis thaliana* precursors, obtained from ProteomXchange (dataset PXD012710, *Arabidopsis* proteome spectral library, “Arabidopsis_Library_TripleTOF5600_Spectronaut.xls”) as negative controls. Peptides that matched to both, the UniProt^45^ Human canonical proteome (3AUP000005640) as well as the UniProt *Arabidopsis thaliana* canonical proteome (3AUP000005648) were removed from the library. The spectra and retention times in the merged Human/*Arabidopsis thaliana* library were replaced with in silico predicted values using the deep learning-based prediction integrated in DIA-NN. The empirical FDR was estimated as previously described ^23^. In short, the empirical FDR is the ratio of *Arabidopsis thaliana* precursors identified and Human precursors identified multiplied by the ratio of Human precursors and *Arabidopsis thaliana* precursors in the library (only precursors ranging from m/z 400 to m/z 900 were considered).

## Acknowledgments

This work was supported by the Francis Crick Institute which receives its core funding from Cancer Research UK (FC001134), the UK Medical Research Council (FC001134), and the Wellcome Trust (FC001134), and received specific funding from the BBSRC (BB/N015215/1 and BB/N015282/1), as well as a Crick Idea to Innovation (i2i) initiative (Grant Ref 10658) as well as the Crick LifeArc (Project 1290305).

## Data availability

The generated data have been uploaded to Mendeley data (https://data.mendeley.com/datasets/gghrtzr69d/draft?a=63233295-4a34-4706-88a6-a9807a83d036); previously published data were also used for the benchmarks (PXD016662).

## Competing interest

N.B, G.I., F.W and S.T. work for SCIEX

---

**Figure S1:**
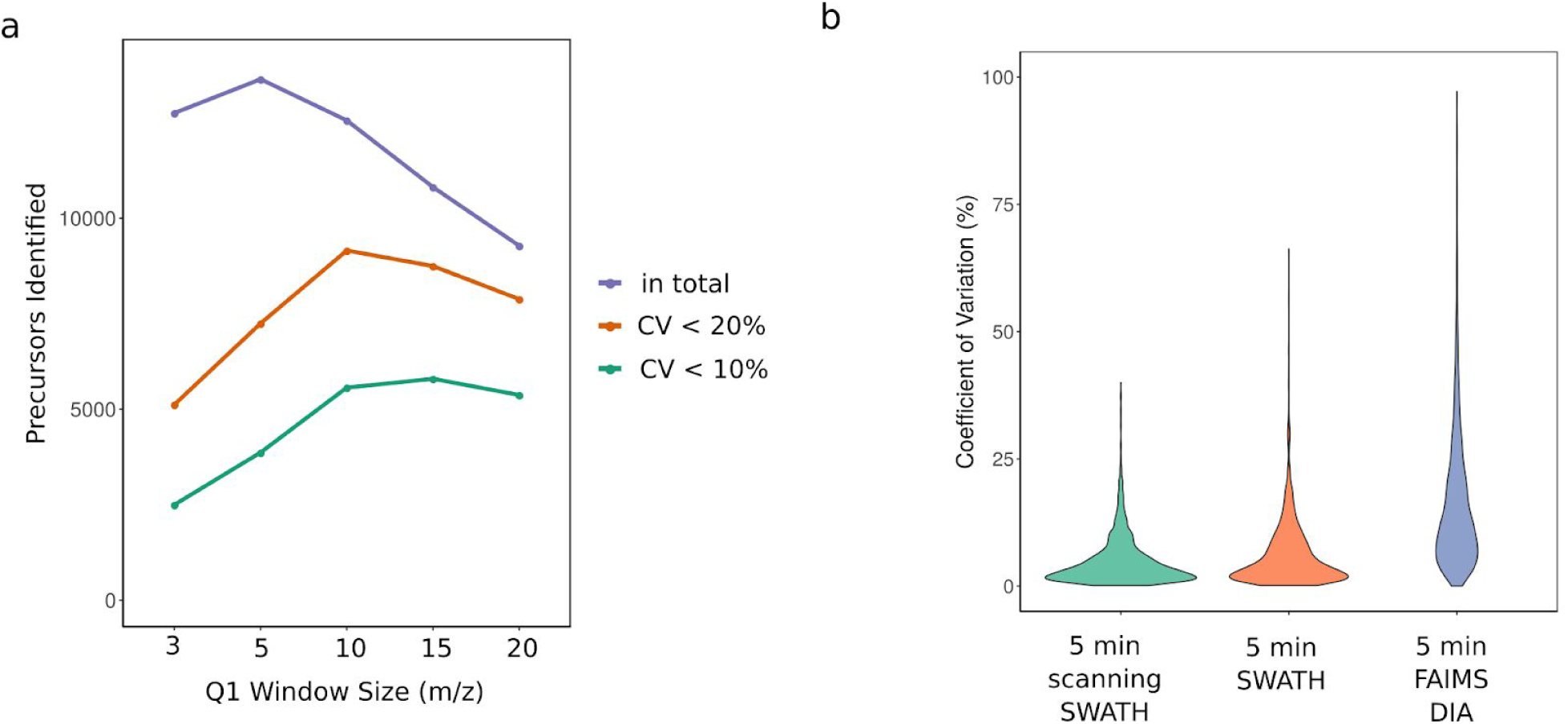
**a.** Comparison of different scanning SWATH precursor isolation window sizes (3 m/z, 5 m/z, 10 m/z and 20 m/z). 5 µg yeast digests was injected, the duty cycle was 0.5 seconds, the precursor range m/z 400 to m/z 900 and the gradient length was 5 minutes. Total number of precursor identifications (1 % FDR) as well as precursors quantified with less than 10 % and 20 % coefficient of variation (CV) in triplicate injections are shown. **b**. Protein group CV values from triplicate injections of cell lysates are compared between acquisition schemes and platforms. Scanning SWATH (10 m/z window) is compared to conventional stepped SWATH (both acquired on a TripleTOF 6600 coupled to an Agilent Infinity II) and DIA on an Orbitrap Exploris 480 with a FAIMS interface coupled to an Evosep One (PXD016662)^19^. In order to make the data comparable between platforms, protein groups were only considered when quantified on each platform (2172 protein groups).

**Table S1.** Lower and upper m/z limits of the SWATH precursor selection windows used.

| Lower m/z limit | Upperr m/z limit | CE spread |
| --- | --- | --- |
| 399.5 | 438.3 | 5 |
| 437.3 | 470.4 | 5 |
| 469.4 | 499.9 | 5 |
| 498.9 | 528 | 5 |
| 527 | 553.8 | 5 |
| 552.8 | 578.8 | 5 |
| 577.8 | 602.9 | 5 |
| 601.9 | 626.5 | 5 |
| 625.5 | 650.2 | 5 |
| 649.2 | 673.4 | 5 |
| 672.4 | 696.1 | 5 |
| 695.1 | 720.2 | 5 |
| 719.2 | 747.4 | 5 |
| 746.4 | 777.3 | 5 |
| 776.3 | 810.3 | 5 |
| 809.3 | 850.9 | 5 |
| 849.9 | 899.5 | 5 |

**Table S2:** Scanning SWATH MS duty cycles for different gradient length.

| Gradient length (minutes) | Duty cycle (secs) | Window size |
| --- | --- | --- |
| 0.5 | 0.28 | 10da |
| 1 | 0.31 | 10da |
| 3 | 0.41 | 10da |
| 5 | 0.52 | 10da |

**Table S3:** Lower and upper m/z limits of the SWATH precursor selection windows used for the different gradient length.

| 30 seconds gradient length |  |
| --- | --- |
| 399.5 | 435.2 |
| 434.2 | 469.8 |
| 468.8 | 505.6 |
| 504.6 | 544.1 |
| 543.1 | 585.3 |
| 584.3 | 629.8 |
| 628.8 | 679.9 |
| 678.9 | 737.1 |
| 736.1 | 806.9 |
| 805.9 | 899.9 |
| 1minute gradient length |  |
| 399.5 | 429.7 |
| 428.7 | 458.3 |
| 457.3 | 487.4 |
| 486.4 | 518.2 |
| 517.2 | 550.6 |
| 549.6 | 585.3 |
| 584.3 | 622.1 |
| 621.1 | 662.8 |
| 661.8 | 707.4 |
| 706.4 | 759.1 |
| 758.1 | 819.6 |
| 818.6 | 899.9 |
| 3minute gradient length |  |
| 399.5 | 423.6 |
| 422.6 | 446.7 |
| 445.7 | 469.8 |
| 468.8 | 493.5 |
| 492.5 | 518.2 |
| 517.2 | 544.1 |
| 543.1 | 571.5 |
| 570.5 | 599.6 |
| 598.6 | 629.8 |
| 628.8 | 662.8 |
| 661.8 | 698 |
| 697 | 737.1 |
| 736.1 | 782.2 |
| 781.2 | 833.3 |
| 832.3 | 899.9 |
| 5minute gradient length |  |
| 399.5 | 419.8 |
| 418.8 | 439 |
| 438 | 458.3 |
| 457.3 | 477.5 |
| 476.5 | 497.3 |
| 496.3 | 518.2 |
| 517.2 | 539.7 |
| 538.7 | 562.2 |
| 561.2 | 585.3 |
| 584.3 | 609.5 |
| 608.5 | 635.3 |
| 634.3 | 662.8 |
| 661.8 | 692 |
| 691 | 723.3 |
| 722.3 | 759.1 |
| 758.1 | 798.7 |
| 797.7 | 843.8 |
| 842.8 | 899.9 |

**Table S4:** MS-duty cycles for different gradient length.

| Gradient length (minutes) | Duty cycle (secs) | Number of windows |
| --- | --- | --- |
| 0.5 | 0.31 | 10 |
| 1 | 0.36 | 12 |
| 3 | 0.43 | 15 |
| 5 | 0.51 | 18 |

## Notes

### Competing Interest Statement

N.B, G.I., F.W and S.T. are employees of SCIEX

### Summary of Updates

This version of the manuscript has been revised with improved and expanded results.

